# SpyChIP identifies cell type-specific transcription factor occupancy from complex tissues

**DOI:** 10.1101/2022.03.25.485871

**Authors:** Siqian Feng, Richard S. Mann

## Abstract

Chromatin immunoprecipitation (ChIP) is an important technique for characterizing protein-DNA binding *in vivo*. One drawback of ChIP based techniques is the lack of cell type-specificity when profiling complex tissues. To overcome this limitation, we developed SpyChIP to identify cell type-specific transcription factor (TF) binding sites in native physiological contexts without tissue dissociation or nuclei sorting. SpyChIP takes advantage of a specific covalent isopeptide bond that rapidly forms between the 15 amino acid SpyTag and the 17 kD protein SpyCatcher. In SpyChIP, the target TF is fused with SpyTag by genome engineering, and an epitope tagged SpyCatcher is expressed in cell populations of interest, where it covalently binds to SpyTag-TF. Cell type-specific ChIP is obtained by immunoprecipitating chromatin prepared from whole tissues using antibodies directed against the epitope-tagged SpyCatcher. Using SpyChIP, we identified the genome-wide binding profiles of the Hox protein Ubx in two distinct cell types of the *Drosophila* haltere disc. Our results revealed extensive region-specific Ubx-DNA binding events, highlighting the significance of cell type-specific ChIP and the limitations of whole tissue ChIP approaches. Analysis of Ubx::SpyChIP results provided novel insights into the relationship between chromatin accessibility and Ubx-DNA binding, as well as different mechanisms Ubx employs to regulate its downstream *cis*-regulatory modules (CRMs). In addition to SpyChIP, we suggest that SpyTag-SpyCatcher technology, as well as other covalent interaction peptide pairs, has many potential *in vivo* applications that were previously unachievable.

## Introduction

Chromatin immunoprecipitation followed by high throughput sequencing (ChIP-seq) has been an important technique to query *in vivo* genome-wide binding profiles of transcription factors (TFs) and chromatin modifications (1). However, when assayed in whole tissues, ChIP-seq reports a mixture of TF-DNA binding signatures present in multiple cell types, making it difficult to decern a TF’s cell type-specific functions. Several strategies have been developed to obtain cell type-specific TF-DNA occupancy information. Cell type-specific overexpression of tagged TFs is not an ideal solution, because non-physiological levels or non-native spatial and/or temporal expression patterns can result in false positive or false negative binding. An alternative is to sort crosslinked nuclei from dissociated tissues (2), but dissociation remains a significant technical challenge for many tissues, and the low yield of sorting makes this strategy only feasible for tissues that can be obtained in large quantity. Targeted DamID (TaDa), which depends on cell type-specific expression of very low levels DNA adenine methyltransferase (Dam)-TF fusions, represents another powerful approach (3). However, it can be challenging to accurately control the levels of the TF-Dam fusions, and DamID-based methods have the potential to mark a mixture of past and present TF binding events, compromising the temporal resolution of the results that may be important when characterizing actively developing tissues.

To overcome the limitations of the current techniques, we developed a method based on SpyTag-SpyCatcher technology (4) that we call SpyChIP. Previous *in vitro* work demonstrated that the 15 amino acid SpyTag peptide spontaneously and rapidly forms a covalent isopeptide bond with a specific binding partner, a 17 kD protein named SpyCatcher (4). We reasoned that if SpyTag and SpyCatcher were also able to form a covalent bond in nuclei, a TF fused with SpyTag could be covalently linked to epitope tagged spyCatcher expressed specifically in the target cell type. ChIP against the epitope on spyCatcher would decode cell type-specific TF-DNA occupancy without tissue dissociation and nuclei sorting (Fig. 1A). Indeed, applying SpyChIP to the *Drosophila* Hox protein Ubx verified this approach and revealed many cell type-specific Ubx-DNA binding events in the haltere imaginal disc.

**Fig. 1.**
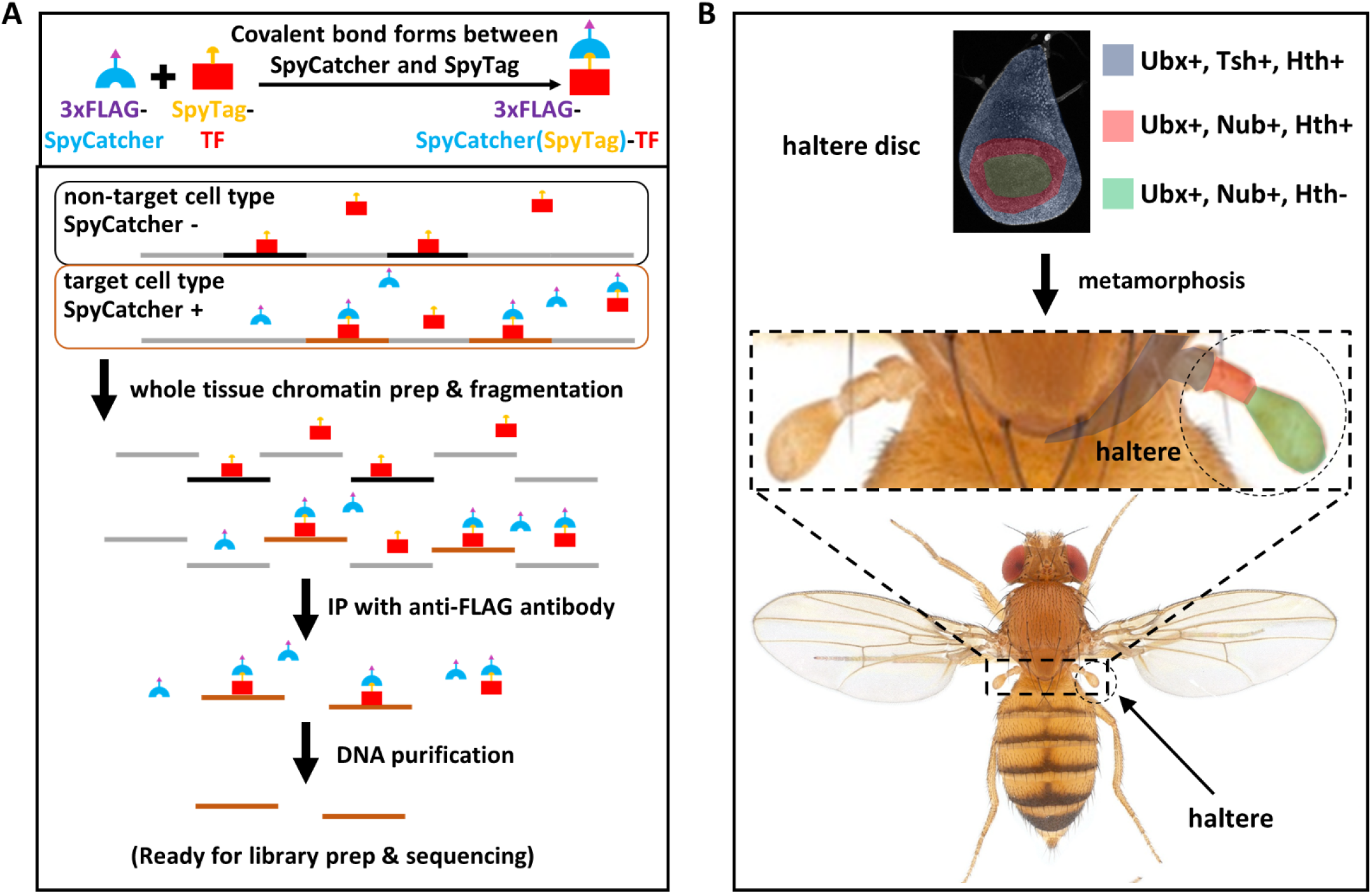
Overview of SpyChIP strategy and haltere development. **A.** A TF of interest is tagged with SpyTag by genome engineering. Upon cell-type specific expression of 3xFLAG-SpyCatcher, a covalent bond is formed between SpyTag and SpyCatcher, allowing chromatin bound by the TF to be immunoprecipitated using antibody against the 3xFLAG epitope on SpyCatcher. **B.** Schematic of the development from larval haltere imaginal disc to adult T3 segment. During metamorphosis, the center of the haltere disc everts and becomes the distal haltere. Ubx is expressed in the entire haltere disc. The expression domains of Tsh, Nub and Hth in the haltere disc are labeled, and the corresponding adult structures are indicated by the same colors.

## Results

### SpyTag and SpyCatcher form a covalent isopeptide bond *in vivo*

We first tested whether SpyTag and SpyCatcher form a covalent isopeptide bond *in vivo*. In the nuclei of *Drosophila* embryos, we co-expressed 3xFLAG-SpyCatcher with GFP that was tagged with SpyTag at either the N- or C-terminus, and the V5 tag at the other end. Western blot against the 3xFLAG tag and the V5 tag was performed to follow SpyCatcher and GFP respectively. Consistent with previous *in vitro* results, we detected the formation of a larger molecular weight protein that is roughly the predicted size of SpyCatcher fused to GFP (Fig. 2A), indicating successful covalent bond formation in *Drosophila* nuclei.

**Fig. 2.**
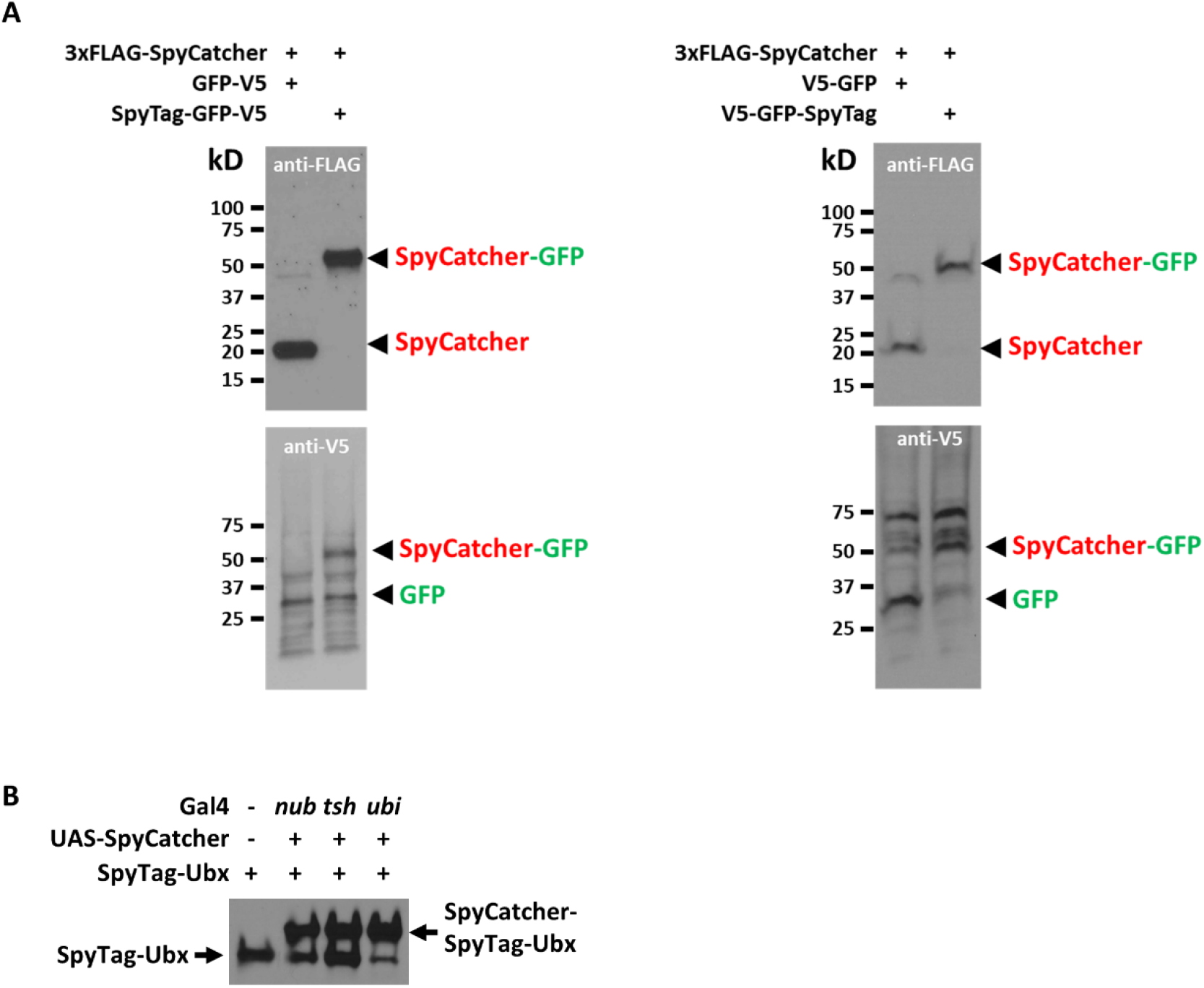
SpyTag and SpyCatcher form covalent isopeptide bond *in vivo*. **A.** Western blot analysis of total embryo lysates using anti-FLAG antibody and anti-V5 antibody. The predicted molecular weights of relevant proteins are shown below. The embryos were F1 embryos from the following crosses: Left: SpyTag at N-terminus of GFP: *En-Gal4/(CyO); MKRS/TM6B* males crossed to *attP40-UAS-3xFLAG-NLS-SpyCatcher; attP2-UAS-(SpyTag)-GFP-V5* females. Right: SpyTag at C-terminus of GFP: *En-Gal4/(CyO)* females crossed to *attP40-UAS-3xFLAG-NLS-SpyCatcher; attP2-UAS-V5-GFP-(SpyTag)* males. In both cases, only GFP that is tagged with SpyTag shifts to a higher molecular weight after expression of 3xFLAG-SpyCatcher. **B.** Anti-Ubx western blot analysis of whole haltere discs. The genotypes of the lanes from left to right are: 1) *SpyTag-Ubx/SpyTag-Ubx*, 2) *nub-Gal4/+; UAS-SpyCatcher, SpyTag-Ubx/SpyTag-Ubx*, 3) *tsh-Gal4/+; UAS-SpyCatcher, SpyTag-Ubx/SpyTag-Ubx*, and 4) *ubi-Gal4/+; UAS-SpyCatcher, SpyTag-Ubx/SpyTag-Ubx*. Depending on the driver, different amounts of SpyTag-Ubx are shifted to a higher molecular weight upon co-expression with SpyCatcher.

We next piloted SpyChIP by characterizing the occupancy of the Hox protein Ubx (Ultrabithorax) in different cell types in *Drosophila* haltere imaginal discs. Ubx is a selector TF that determines the identity of the 3^rd^ thoracic (T3) and 1^st^ abdominal (A1) segments (5). We probed the genome-wide binding of Ubx in the *Drosophila* haltere imaginal disc, which gives rise to the dorsal T3 segment of the adult fly, including the haltere, an appendage critical for flight. Mutations in *Ubx* result in the famous four-winged *bithorax* homeotic transformation, in which the haltere-bearing T3 segment of the adult is transformed into a second copy of the wing-bearing T2 segment (Fig. 1B) (5). During wild type metamorphosis, the center of the haltere imaginal disc gives rise to most of the haltere appendage, while the periphery of the disc gives rise to the dorsal T3 body wall and the proximal haltere structures (Fig. 1B) (6).

We fused the SpyTag to the N-terminus of Ubx at the endogenous *Ubx* locus in a scarless manner (Fig S1 and Methods), and expressed 3xFLAG-SpyCatcher with 2 cell type-specific Gal4 drivers: *tsh-Gal4*, active in the proximal haltere disc, and *nub-Gal4*, expressed in the distal haltere disc. We also used the ubiquitous driver *ubi-Gal4*, which should mimic a standard whole tissue ChIP experiment (Fig. 1B and Fig. S2). Western blotting with an anti-Ubx antibody showed that the apparent molecular weight of SpyTag-Ubx increased when SpyCatcher was expressed by all three drivers, and that the increase in size was consistent with the molecular weight of 3XFLAG-SpyCatcher (Fig. 2B). When *ubi-Gal4* was used to express SpyCatcher, most of the endogenous Ubx shifted to the larger molecular weight (Fig. 2B), indicating efficient covalent bond formation between SpyCatcher and SpyTag-Ubx *in vivo* in *Drosophila* nuclei. As expected, when SpyCatcher was expressed with the other two drivers, less Ubx was shifted to the larger size, consistent with their more limited expression domains within the haltere disc.

### SpyChIP faithfully captures TF-DNA occupancy

ChIP-seq experiments were then performed when 3xFLAG-SpyCatcher was expressed by each of the three Gal4 drivers, using chromatin prepared from whole haltere discs and anti-FLAG antibody. All Ubx::SpyChIP replicates revealed thousands of peaks, consistent with successful ChIP experiments. To assess how well SpyChIP works, we compared *ubi-Gal4*>Ubx::SpyChIP results with 2 independent whole haltere disc Ubx ChIP datasets. One such dataset was generated by using the same anti-FLAG antibody as we used in all Ubx::SpyChIP experiments to profile Ubx binding in 3xFLAG-Ubx flies, which was previously created by inserting the 3xFLAG tag into the endogenous *Ubx* locus in a scarless manner (7). The other whole disc Ubx ChIP dataset was obtained by probing wild type flies using anti-Ubx antibody (8). The average enrichment of sequencing tags in all called peaks relative to a random set of genomic regions can be used as an approximation of a ChIP’s signal-to-noise ratio. We found that this enrichment is slightly higher for Ubx ChIP with anti-FLAG antibody than with anti-Ubx antibody (Fig. S3A). All Ubx::SpyChIP experiments have similar enrichment, which is essentially the same as the enrichment of Ubx ChIP with anti-Ubx antibody, but is slightly lower than anti-FLAG Ubx ChIP (Fig. S3A). We conclude that overall, the signal-to-noise ratio of SpyChIP is comparable to that of standard ChIP experiments.

In addition, pair-wise comparisons between *ubi-Gal4>Ubx::SpyChIP* and both whole haltere disc Ubx ChIPs show good agreement (Fig. 3A and Fig. S3B). The correlation between *ubi-Gal4*>Ubx::SpyChIP and a standard Ubx ChIP is similar to the correlation between two Ubx ChIP biological replicates (Fig. S3B), indicating that SpyChIP faithfully captures genome-wide Ubx occupancy.

**Fig. 3.**
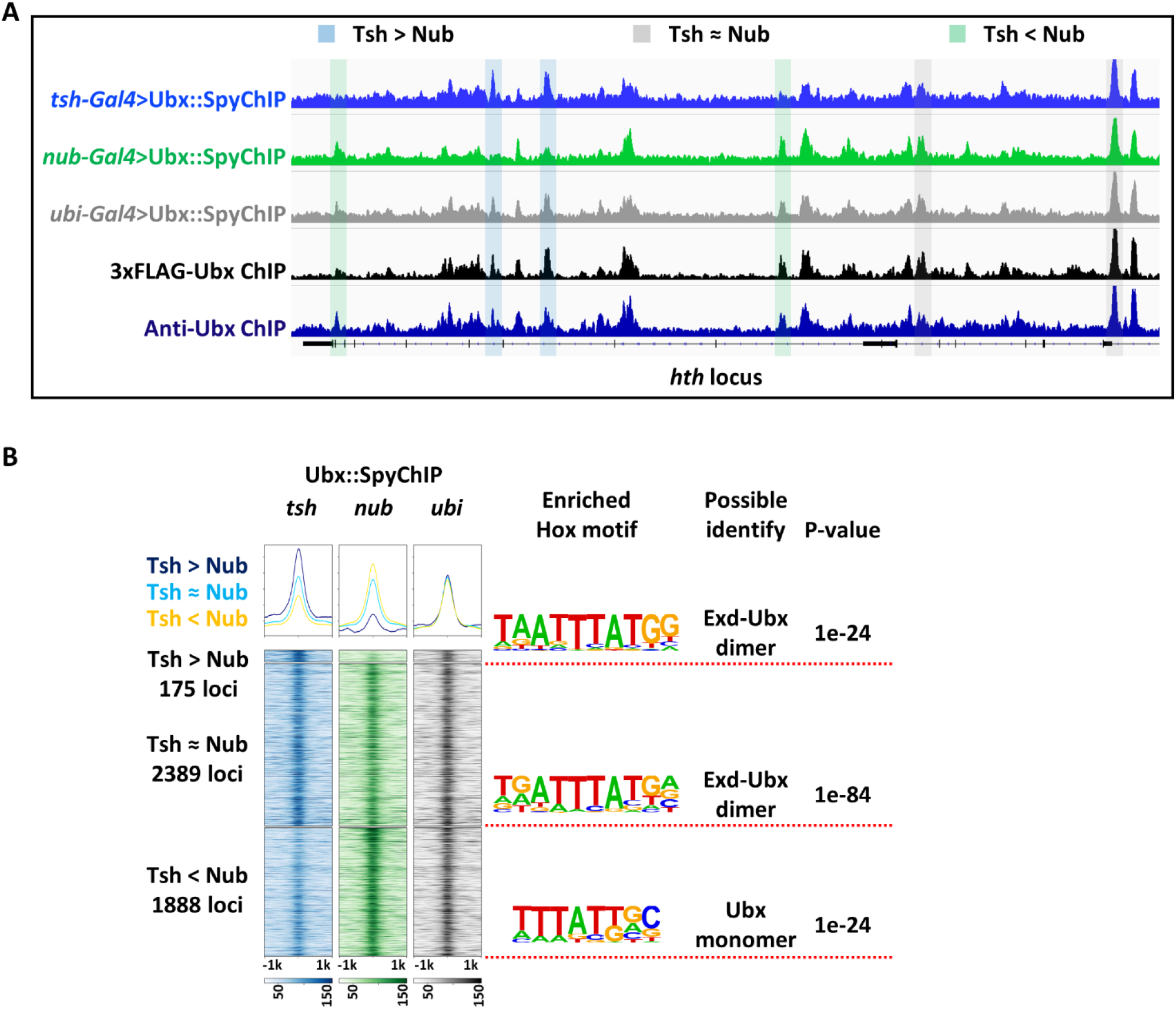
SpyChIP identifies genome-wide and cell type-specific TF binding events. **A.** SpyChIP results at the *hth* locus, which was chosen as an example. Examples of different classes of peaks are color coded: blue: Tsh Ubx::SpyChIP > Nub Ubx::SpyChIP, grey: Tsh Ubx::SpyChIP ≈ Nub Ubx::SpyChIP, and green: Tsh Ubx::SpyChIP < Nub Ubx::SpyChIP. Three SpyChIP tracks and two independent whole haltere disc Ubx ChIP tracks are shown. The 3xFLAG-Ubx (7) ChIP used the same anti-FLAG antibody as in all SpyChIP experiments. For comparison, the anti-Ubx ChIP track used an antibody directed against Ubx (8). **B.** Heatmaps and histograms of Tsh > Nub, Tsh ≈ Nub and Tsh < Nub Ubx::SpyChIP loci plotted for *tsh-Gal4*>Ubx::SpyChIP, *nub-Gal4*>Ubx::SpyChIP and *ubi-Gal4*>Ubx::SpyChIP. Hox-related motifs significantly enriched in each class of loci are indicated. For a complete list of enriched motifs, see Fig. S4.

We considered the possibility that, when SpyCatcher is expressed with *nub-Gal4* or *tsh-Gal4*, there may be a large excess of SpyCatcher compared to SpyTag-Ubx. Such an excess could result in a pool of unbound SpyCatcher that, during chromatin preparation and immunoprecipitation, might bind to SpyTag-Ubx from cells outside the domain targeted by Gal4, thus potentially compromising specificity. To limit this from happening, an excess of synthetic SpyTag peptide was added to quench unoccupied SpyCatcher in all experiments except for *nub-Gal4*>Ubx::SpyChIP replicate 1, which allowed us to assess the effect of quenching. The comparison between *nub-Gal4*>Ubx::SpyChIP replicates with or without quenching did not reveal significant differences (Fig. S3C). This could mean that an excess of SpyCatcher does not decrease the specificity of SpyChIP or, in this case, it could be due to the fact that the endogenous Ubx levels are sufficiently high in Nub+ cells (Delker et al., 2019) so that there is not an excess of unbound SpyCatcher.

### SpyChIP identifies cell type-specific TF-DNA binding events

We next inspected Ubx::SpyChIP results genome-wide. Peaks shared between Tsh+ and Nub+ cells, as well as those specific to each cell type, could be readily identified (Fig. 3A). Genome-wide comparison between *tsh-Gal4*>Ubx::SpyChIP and *nub-Gal4*>Ubx::SpyChIP results identified 175 and 1888 Ubx binding events that are specific to either the Tsh+ domain or Nub+ domain, respectively. In addition, there are 2389 binding events that are shared by both datasets (Fig. 3B). The significant asymmetry in the numbers of Tsh+ and Nub+ cell-specific Ubx binding events is surprising, but is consistent with the observation that for both the wing and haltere discs, several fold more differentially accessible loci were observed in Nub+ cells than in Tsh+ cells (8).

Ubx can bind to DNA either as a monomer or as a heterodimer with its cofactor Extradenticle (Exd), and the ubiquitous Exd protein is only nuclear and available as a Hox cofactor when another protein, Homothorax (Hth), is present (9). In the haltere disc, Hth is expressed in all Tsh+ cells and some Nub+ cells (8) (Fig. 1B). Consistent with the large number of Nub+, Hth-cells, a Ubx monomer motif is enriched in Nub+ cell-specific Ubx-bound peaks. In contrast, an Exd-Ubx heterodimer motif is enriched in Tsh+ cell-specific Ubx binding events, as well as in peaks shared by the two cell types (Fig. 3B and Fig. S4B). As expected, both types of Ubx motifs are enriched in the *ubi-Gal4>*Ubx::SpyChIP peaks (Fig. S4A). These results are consistent with previous results showing that Ubx binds with or without cofactors, depending on the region of the haltere disc (8), and demonstrate that SpyChIP is able to capture cell type-specific TF-DNA binding events.

### The role of cell type-specific Ubx binding

Recently, Loker *et. al*. characterized the genome-wide chromatin accessibility in Tsh+ and Nub+ cells of the haltere and the serially homologous wing imaginal discs (8). Given the cell type-specific Ubx binding data described here, we asked if there is any correlation between cell type-specific chromatin accessibility and cell type-specific Ubx binding. Notably, sites in the haltere that have Tsh>Nub Ubx binding also tend to be more accessible in Tsh+ cells compared to Nub+ cells, not only in the haltere disc, but also in the wing disc (Fig. 4A and 4B). Since Ubx is expressed in the haltere disc but not in the wing disc, this pattern suggests that the 175 Tsh>Nub Ubx binding sites gain accessibility in Tsh+ cells by a mechanism that is independent of Ubx binding. Similarly, many, but not all of the 1888 Nub>Tsh Ubx binding sites have biased accessibility in Nub+ cells compared to Tsh+ cells, in both the haltere and wing (Fig. 4C and 4D).

**Fig. 4.**
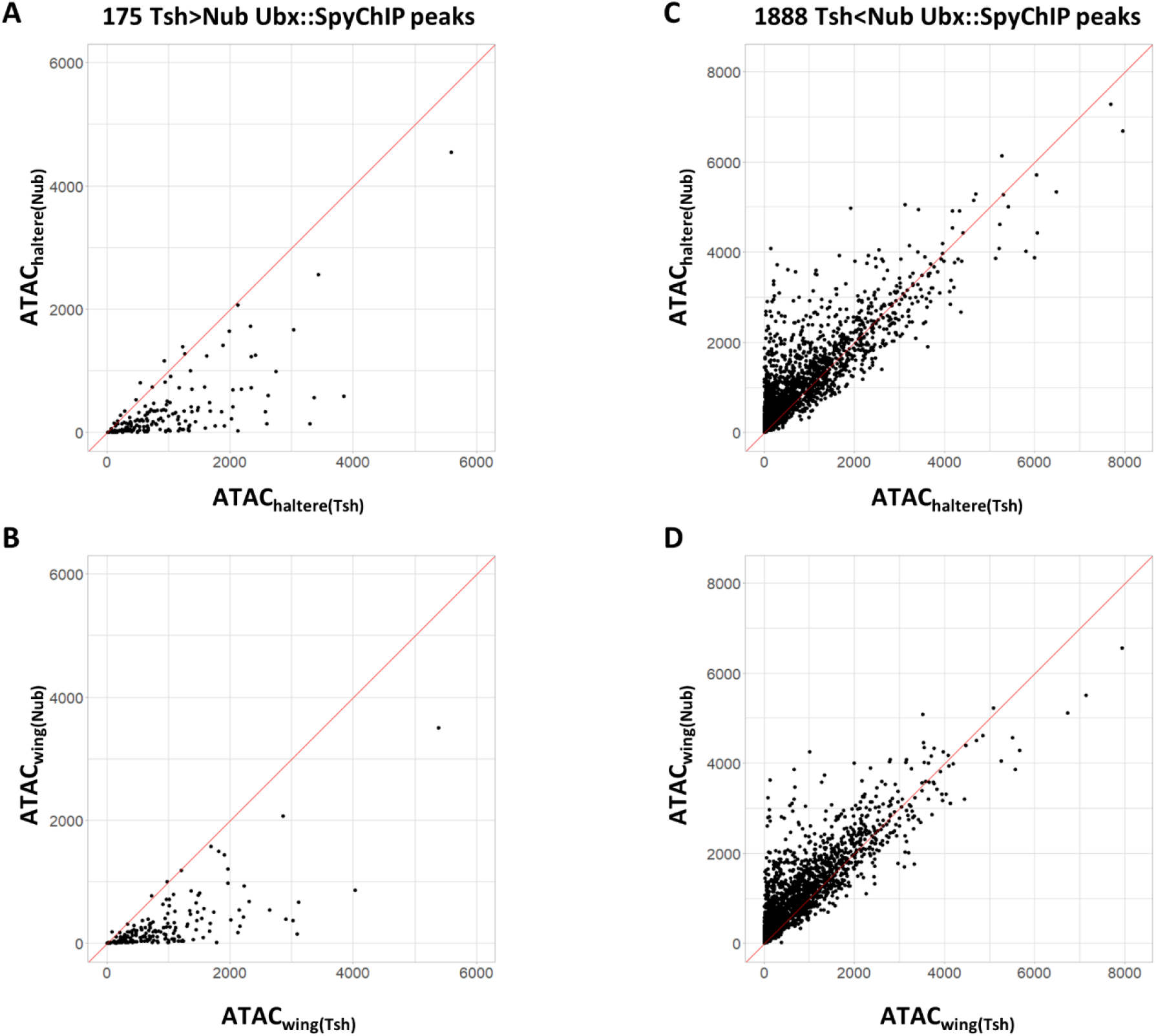
Relationship between chromatin accessibility and TF-DNA binding revealed by SpyChIP. Scatter plots comparing chromatin accessibility of Tsh+ and Nub+ cells in 175 Tsh > Nub Ubx::SpyChIP peaks (**A** and **B**), or in 1888 Tsh < Nub Ubx::SpyChIP peaks (**C** and **D**). The Tsh+ vs. Nub+ cells were compared in both the haltere disc (**A** and **C**) and the wing disc (**B** and **D**). Chromatin accessibility data are from (8).

Finally, we inspected Ubx::SpyChIP patterns at selected Ubx downstream *cis*-regulatory modules (CRMs). For simplicity, we focused on CRMs that only require Ubx function in Nub+ cells and also have Ubx ChIP peaks from whole haltere disc experiments, suggesting that they are direct Ubx targets. We included in our analysis *sal1.1* (10) and *kn01* (11), as well as 4 additional CRMs recently identified by Loker *et. al.* based on their differential accessibility in haltere Nub+ cells compared to wing Nub+ cells (8). Ubx acts as either an activator or a repressor of each CRM (Fig. 5). Among the 6 selected CRMs, 4 have Ubx binding only in Nub+ cells, while the other 2 have Ubx binding in both Tsh+ and Nub+ cells. These patterns of binding and regulation are consistent with the existence of multiple modes of Ubx regulation. For example, for CRM Rep-6, which is activated by Ubx in Nub+ cells, Ubx binding is observed in both Tsh+ and Nub+ cells and is apparently not sufficient for activation of this CRM. In contrast, Ubx only binds to CRM Rep-7 in Nub+ cells, where it also acts as an activator, raising the possibility that the absence of Ubx binding in Tsh+ cells is important for CRM activation only in Nub+ cells.

**Fig. 5.**
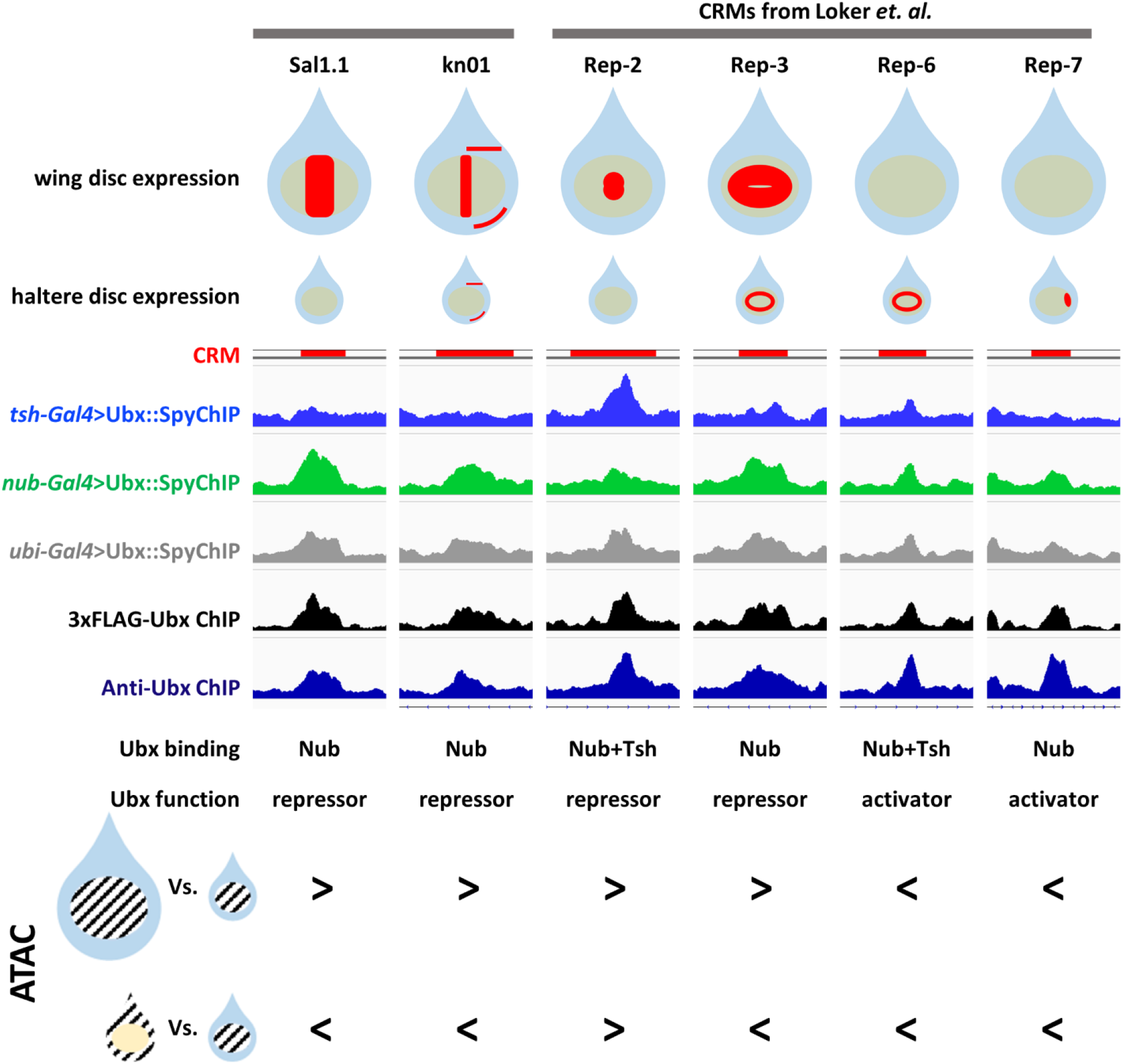
SpyChIP reveals distinct Ubx regulatory strategies. Summary of Ubx binding, expression patterns and chromatin accessibility for selected Ubx-targeted CRMs. These CRMs were chosen because they bind Ubx and have been shown to require *Ubx* function, either as a repressor or activator as indicated, in Nub+ cells (8). The top two rows are schematics of the CRM expression patterns in wing and haltere discs. Light blue and yellow colors mark the Tsh+ and Nub+ cells, respectively; red indicates CRM activity. Below are five genome browser views showing the Ubx::SpyChIP signals and whole disc Ubx ChIP signals, relative to the location of the CRMs (red bars). The bottom two rows compare the patterns of chromatin accessibility (8) between the Nub+ cells of the wing vs haltere (top row) and the Tsh+ vs. Nub+ cells in the haltere (bottom row), for each CRM. Note that Ubx activity as a repressor or activator correlates with less or more accessibility, respectively, in haltere Nub+ cells compared to wing Nub+ cells. Also notable is that the four examples that have Tsh < Nub Ubx binding also have Tsh < Nub chromatin accessibility. In contrast, in the two cases where Ubx binding is observed in both Nub+ and Tsh+ cells, there is no correlation with accessibility differences.

## Discussion

Characterizing cell type-specific binding is critical for understanding a TF’s *in vivo* functions. The SpyChIP technique we describe in this study overcomes several major limitations of existing approaches. Because SpyChIP does not depend on tissue dissociation or nuclei sorting, it is especially suitable for tissues with limited availability or those that are difficult to dissociate. Contrary to the lower temporal resolution associated with DamID based techniques, the temporal resolution of SpyChIP has as high a temporal resolution as standard ChIP and is therefore desirable in analyzing tissues undergoing dynamic rearrangements. We demonstrated the efficacy of SpyChIP by successfully obtaining cell type-specific Ubx ChIP results from the *Drosophila* haltere discs. These tiny tissues must be manually dissected and are therefore difficult to obtain in large quantity. Imaginal discs also undergo rapid cellular rearrangements during metamorphosis. In fact, to our knowledge, before our study, no cell type-specific TF-DNA occupancy results have been reported from any *Drosophila* imaginal discs. The covalent bond between Spytag and SpyCatcher is robust to diverse conditions such as temperature and pH (4), thus SpyChIP is likely to be applicable in most tissues and in most organisms. If the target cell type represents a very small fraction in the complex tissue, SpyChIP may be combined with crude cell/nuclei sorting to partially enrich the target cells. In SpyChIP, cell type-specificity is genetically encoded, it is thus not necessary to obtain a highly pure cell population by sorting, which is usually associated with lower yields.

Although a positive correlation is often observed between differential chromatin accessibility and differential transcription factor binding, it is usually difficult to deduce the cause versus the consequence. With the aid of Ubx::SpyChIP, we were able to rule out that Tsh>Nub Ubx binding caused Tsh>Nub chromatin accessibility. Conversely, our results suggest that Tsh>Nub chromatin accessibility is permissive for Tsh>Nub Ubx binding pattern. It is generally believed that the same TF, especially a selective TF like Ubx, can regulate its downstream CRMs using different modes of action. However, it is not easy to demonstrate diverse mechanisms. Our Ubx::SpyChIP results show that Ubx binding is not always sufficient for CRM activation, suggesting the presence of multiple mechanisms that act in a CRM-specific manner.

Finally, we suggest that the SpyTag/SpyCatcher technology has the potential for many additional *in vivo* applications beyond SpyChIP. We envision that the covalent interaction between Spytag and SpyCatcher can be combined with a variety of other techniques, such as HiChIP (12) and bioID (13), to achieve cell type-specificity without dissociation or cell/nucleus sorting. Moreover, once a factor has been fused with SpyTag by genome modification, it can be easily tagged with any peptide of interest, such as different epitopes, fluorescent proteins, or enzymes. Also noteworthy is that SpyTag and SpyCatcher are not the only pair of peptides that form a covalent bond when they interact: other orthogonal pairs have been reported to for covalent bonds *in vitro* (14). Therefore, there are many possibilities of *in vivo* applications of these covalent interacting peptide pairs.

## Materials and Methods

### New fly strains

_All plasmid were generated by standard procedures, and transgenic flies were generated by integrating the plasmids into selected attP sites via phiC31 integrase mediated site-specific recombination.

The scarless *SpyTag-Ubx* allele was generated using a method we previously described (7). Briefly, a fragment of *Ubx* genomic DNA containing the *SpyTag* inserted at the N-terminal end of the *Ubx* ORF was integrated into the endogenous *Ubx* locus by phiC31 integrase mediated site-specific recombination. Double-stranded DNA breaks were then introduced to stimulate homologous recombination and repair the endogenous *Ubx* to the final scarless *SpyTag-Ubx* allele. The landing site for site-specific recombination in the *Ubx* locus has been described in detail, and the donor plasmid was generated similarly as before (7). The *SpyTag* sequence was inserted by overlapping extension PCR. Multiple independent *SpyTag-Ubx* alleles were generated, verified by southern blotting, and fully sequenced to make sure there were no unwanted mutations. Southern blotting was performed using DIG High Prime DNA Labeling and Detection Starter Kit II (Roche 11585614910) and DIG Wash and Block Buffer Set (Roche 11585762001) according to manufacturer’s instructions. The *Ubx* 5’ and *Ubx* 3’ probes were described before (7). DNA Molecular Weight Marker II, DIG-labeled (Roche 11218590910) was used as the marker.

### Western blotting

Western blotting was performed using standard procedure. For embryo samples, embryos from desired crosses were collected overnight at 25°C, and transferred to a 1ml Wheaton homogenizer (not dechorionated). An appropriate volume of 4xSDS-PAGE loading dye (with 10% β-mercaptoethanol) was added (100ul of the loading dye per ~10ul of settled embryos), and the embryos were completely homogenized. The homogenized materials were then transferred to 1.5ml tubes. For each haltere disc sample, 35-55 discs were dissected in PBS+1% BSA on ice, and transferred to a 1.5ml tube containing 0.5ml of PBS+1% BSA. The supernatant was removed, and 100ul of 4xSDS-PAGE loading dye (with 10% β-mercaptoethanol) was added. The haltere discs were then completely homogenized with a disposable pestle. The homogenized materials were heated at 95°C for 6-7 minutes and chilled on ice. The samples were then spun at room temperature at max speed for 5 minutes, and the supernatant was loaded on SDS-PAGE. After SDS-PAGE, the proteins were transferred to PVDF membrane using routine procedure. The 3xFLAG epitope was detected using anti-FLAG M2-HRP (sigma A8592, 1:10,000), and the V5 epitope was detected using mouse anti-V5 antibody (Invitrogen R96025, 1:5,000) followed by goat anti-mouse IgG-HRP (Jackson ImmunoResearch 115-035-003, 1:25,000), or with rabbit anti-V5 antibody (abcam ab9116, 1:5,000) followed by donkey anti-rabbit IgG-HRP (Jackson ImmunoResearch 711-036-152, 1:5,000). The Ubx protein was detected using monoclonal mouse anti-Ubx FP3.38 (DSHB) at 1:100, followed by the same goat anti-mouse IgG-HRP secondary antibody at 1:10,000. SuperSigna West Pico PLUS Chemiluminescent Substrate (Thermo Scientific 34580) was used as the substrate to visualize the bands.

### Chromatin preparation

The larvae for Ubx::SpyChIP experiments were prepared by crossing *SpyTag-Ubx/(TM6B)* females to *Gal4/(CyO, GFP); attP2-UAS-3xFLAG-NLS-SpyCatcher, SpyTag-Ubx/(TM6B)* males. 3 different *Gal4* lines: *tsh-Gal4, nub-Gal4* and *ubi-Gal4* were used. *TM6B-* and *GFP-* larvae were selected for dissection, and 100 to 150 larvae were dissected for each replicate. Homozygous *3xFLAG-Ubx* (7) larvae were also used for whole haltere disc ChIP experiment. The larvae were pulled apart in PBS and the heat parts were inverted. The inverted heat parts were crosslinked in 10ml of crosslinking solution (10mM HEPES pH8.0, 100mM NaCl, 1mM EDTA pH8.0, 0.5mM EGTA pH8.0, 1% formaldehyde) for 10 minutes at room temperature. After crosslinking, 1ml of 2.5M glycine was added and the samples were hand mixed for 30 seconds. The samples were then washed with 10ml of quenching solution (1xPBS, 125mM glycine, 0.1% Triton X-100) for at least 6 minutes at room temperature, followed by 2 more washes with 10ml of ice-cold buffer A (10mM HEPES pH8.0, 10mM EDTA pH8.0, 0.5mM EGTA pH8.0, 0.25% Triton X-100, with proteinase inhibitor cocktail) at 4°C, 10 minutes each. The gut, salivary glands and fat bodies were then moved from all head parts in buffer A. Next, the samples were washed twice with 10ml of ice-cold buffer B (10mM HEPES pH8.0, 200mM NaCl, 1mM EDTA pH8.0, 0.5mM EGTA pH8.0, 0.01% Triton X-100, with proteinase inhibitor cocktail) at 4°C, 10 minutes each. The haltere discs were dissected from the head parts in buffer B, and were transferred to a 15ml falcon tube. The supernatant was removed, and 0.9ml of buffer C (10mM HEPES pH8.0, 1mM EDTA pH8.0, 0.5mM EGTA pH8.0, 1% Triton X-100, with proteinase inhibitor cocktail) was added. The discs were then sonicated with Branson Sonifier 450 on ice at 15% amplitude for 12 minutes (15 seconds on/30 seconds off). The sonicated samples were spun at max speed at 4°C for 10 minutes, and the supernatant was transferred to new tubes, flash frozen in liquid N_2_, and stored at −80°C until the next step.

The SpyTag stock solution was prepared by dissolving synthetic SpyTag (Genscript custom peptide synthesis service) in water at a concentration of 1mM. For replicates in which synthetic SpyTag was used to quench unoccupied SpyCatcher molecules, SpyTag was used at a final concentration of 10uM in buffer B when taking haltere discs from the head parts, and in buffer C.

### Chromatin immunoprecipitation

ChIP was performed after all chromatin samples were prepared. The chromatin samples were thawed on ice, and to each sample, 1/4 volume of 5x chromatin dilution buffer (50mM Tris-HCl pH8.0, 5mM EDTA pH8.0, 750mM NaCl, 1% Triton X-100) was added to adjust buffer condition, as well as appropriate volume of 100x Halt Protease Inhibitor Cocktail, EDTA-Free (Thermo Scientific 87785). Next, 10 μg of normal mouse IgG was added to each sample for preclearing, and the samples were rotated at 4°C for 1 hour. 40 μl of protein G agarose beads suspension (Roche 11243233001) (settled beads volume 20 μl) was used for each ChIP and preclearing reaction. The beads were washed twice with 1 ml of RIPA buffer (10mM Tris-HCl pH8.0, 1mM EDTA pH8.0, 150mM NaCl, 1% Triton X-100) for 10 minutes each at 4°C with rotation, and were blocked with blocking solution (RIPA + 1.25mg/ml BSA (Sigma A2153) + 0.25mg/ml tRNA (Roche 10109517001)) for at least 1 hour at 4°C with rotation. The chromatin-normal IgG mixtures were added to blocked beads for preclearing, and were rotated at 4°C for 1 hour. The precleared chromatin was separated from beads by centrifugation. 100ul of each precleared chromatin was taken and stored at −80°C as input. 12.5μl of 100 mg/ml BSA, 25μl of 10 mg/ml tRNA, and 10ug of anti-FLAG M2 antibody (Sigma F1804) were added to the rest of precleared chromatin. The samples were rotated at 4°C overnight.

In the next day, the chromatin samples were added to blocked beads, and were rotated at 4°C for 2 hours. The beads were briefly rinsed with RIPA buffer, and were subjected to the following 10-minute washes at 4°C: 2 washes with RIPA buffer, 1 wash with high salt RIPA buffer (10mM Tris-HCl pH8.0, 1mM EDTA pH8.0, 350mM NaCl, 1% Triton X-100), 1 wash with LiCl buffer (10mM Tris-HCl pH8.0, 1mM EDTA pH8.0, 250mM LiCl, 0.1% IGEPAL CA-630), and 1 wash with TE buffer (10mM Tris-HCl, 1mM EDTA, pH8.0, filtered). All rinses washes were performed with 1ml of ice-cold buffer. After the TE wash, the beads were resuspended in 500μl of TE, and the input samples were also adjusted to 500μl with TE buffer. Next, 5μl of 5M NaCl, 12.5μl of 20% SDS, and 10μl of 1mg/ml RNase (Sigma R5503) were added to each ChIP and input sample, and the samples were incubated at 37°C for 30 minutes with rotation. 20μl of 20mg/ml proteinase K (Roche 03115836001) was then added to each sample. The samples were rotated at 55°C for 2 to 3 hours, and then at 65°C overnight for decrosslinking.

In the third day, all ChIP samples were centrifuged at room temperature at max speed for 5 minutes, and the supernatant was transferred to new tubes. 100μl of 3M sodium acetate (pH 5.2) was added to each sample, and the samples were extracted with phenol:chloroform (1:1) and then with chloroform. 1μl of 20mg/ml glycogen (Roche 10901393001) was then added to each sample, and the DNA was purified by isopropanol precipitation. 30μl of 10mM Tris buffer, pH8.0 was used to dissolved the DNA pellet of each sample. The purified DNA was quantified using Qubit dsDNA HS Assay Kit (Thermo Fisher Scientific Q32854)

### ChIP-seq library preparation and sequencing

ChIP-seq libraries were prepared using the NEBNext Ultra™ II DNA Library Prep Kit (NEB E7103) with modifications. 1-2 ng of ChIP DNA and 8-10 ng of input DNA was used as starting materials. No size selection was performed after adaptor ligation, and 11 PCR cycles were performed for all libraries. After PCR amplification, instead of purifying DNA using 0.9x of beads, the following purification protocol was used: 1.8x of beads was used to purify DNA from the PCR reactions. The DNA was eluted in 52ul of elution buffer, and 50ul was transferred to new tubes. The purified DNA was then subjected to size selection (0.65x for first bead addition, and 0.25x for second bead addition). The DNA was then eluted with 17ul of elution buffer, and 15ul was transferred to new tubes. The sizes of the libraries were determined by bioanalyzer, using Bioanalyzer High Sensitivity DNA Analysis (Agilent 5067-4626), and the libraries were quantified by Qubit dsDNA HS Assay Kit (Thermo Fisher Scientific Q32854). The libraries were sequenced using illumina Nextseq 500 sequencer.

### ChIP-seq data analysis

Mapping and peak calling were performed using tools on galaxy.eu. The reads were mapped to *Drosophila* genome build dm6 by bowtie2 (15) using default settings, and peak calling was performed by MACS2 (16) with the following parameters: --nomodel -extsize 200 (all other parameters were default). Differential binding analysis was performed using DiffBind (17), following default procedures. Heatmaps were generated using deeptools2 (18) (also on galaxy.eu), and scatter plots were generated using the R package ggplot2. *de novo* motif searches were performed using homer (19), and all parameters were default except -size 80.

## Acknowledgement

We want to thank Nicolas Gompel for the fly image in Fig. 1B, and we thank all members in the Mann lab for discussions. This study was supported by NIH grants R35 GM118336 to R. S. M.

## Author contributions

S. F. conceived the study, designed the study with input from R. S. M., and performed all the experiments. Both authors analyzed the results and wrote the manuscript.

## References

1. Park PJ (2009) ChIP-seq: advantages and challenges of a maturing technology. Nature Reviews Genetics 10(10):669–680.

2. Bonn S, et al. (2012) Tissue-specific analysis of chromatin state identifies temporal signatures of enhancer activity during embryonic development. Nature Genetics 44(2):148–156.

3. Southall Tony D, et al. (2013) Cell-Type-Specific Profiling of Gene Expression and Chromatin Binding without Cell Isolation: Assaying RNA Pol II Occupancy in Neural Stem Cells. Developmental Cell 26(1):101–112.

4. Zakeri B, et al. (2012) Peptide tag forming a rapid covalent bond to a protein, through engineering a bacterial adhesin. Proceedings of the National Academy of Sciences 109(12):E690.

5. Lewis EB (1963) Genes and Developmental Pathways. American Zoologist 3(1):33–56.

6. Cohen SM (1993) Imaginal disc development. The Development of Drosophila melanogaster., eds Bate M & Martinez-Arias A (Cold Spring Harbor Laboratory Press, New York), Vol 2, pp 747–842.

7. Feng S, Lu S, Grueber WB, & Mann RS (2021) Scarless engineering of the Drosophila genome near any site-specific integration site. Genetics 217(3).

8. Loker R, Sanner JE, & Mann RS (2021) Cell-type-specific Hox regulatory strategies orchestrate tissue identity. Current Biology 31(19):4246–4255.e4244.

9. Merabet S & Mann RS (2016) To Be Specific or Not: The Critical Relationship Between Hox And TALE Proteins. Trends in genetics: TIG 32(6):334–347.

10. Galant R, Walsh CM, & Carroll SB (2002) Hox repression of a target gene: extradenticle-independent, additive action through multiple monomer binding sites. Development (Cambridge, England) 129(13):3115–3126.

11. McKay Daniel J & Lieb Jason D (2013) A Common Set of DNA Regulatory Elements Shapes *Drosophila* Appendages. Developmental Cell 27(3):306–318.

12. Mumbach MR, et al. (2016) HiChIP: efficient and sensitive analysis of protein-directed genome architecture. Nature methods 13(11):919–922.

13. Long S, Brown KM, & Sibley LD (2018) CRISPR-mediated Tagging with BirA Allows Proximity Labeling in Toxoplasma gondii. Bio Protoc 8(6):e2768.

14. Veggiani G, et al. (2016) Programmable polyproteams built using twin peptide superglues. Proceedings of the National Academy of Sciences of the United States of America 113(5):1202–1207.

15. Langmead B & Salzberg SL (2012) Fast gapped-read alignment with Bowtie 2. Nature methods 9(4):357–359.

16. Zhang Y, et al. (2008) Model-based Analysis of ChIP-Seq (MACS). Genome biology 9(9):R137.

17. Ross-Innes CS, et al. (2012) Differential oestrogen receptor binding is associated with clinical outcome in breast cancer. Nature 481:389.

18. Ramírez F, et al. (2016) deepTools2: a next generation web server for deep-sequencing data analysis. Nucleic Acids Research 44(W1):W160–W165.

19. Heinz S, et al. (2010) Simple combinations of lineage-determining transcription factors prime cis-regulatory elements required for macrophage and B cell identities. Mol Cell 38(4):576–589.

