## Supplementary figures and legends for "SpyChIP identifies cell type-specific transcription factor occupancy from complex tissues"

**This PDF file includes:**

Figures S1 to S4
Extended methods

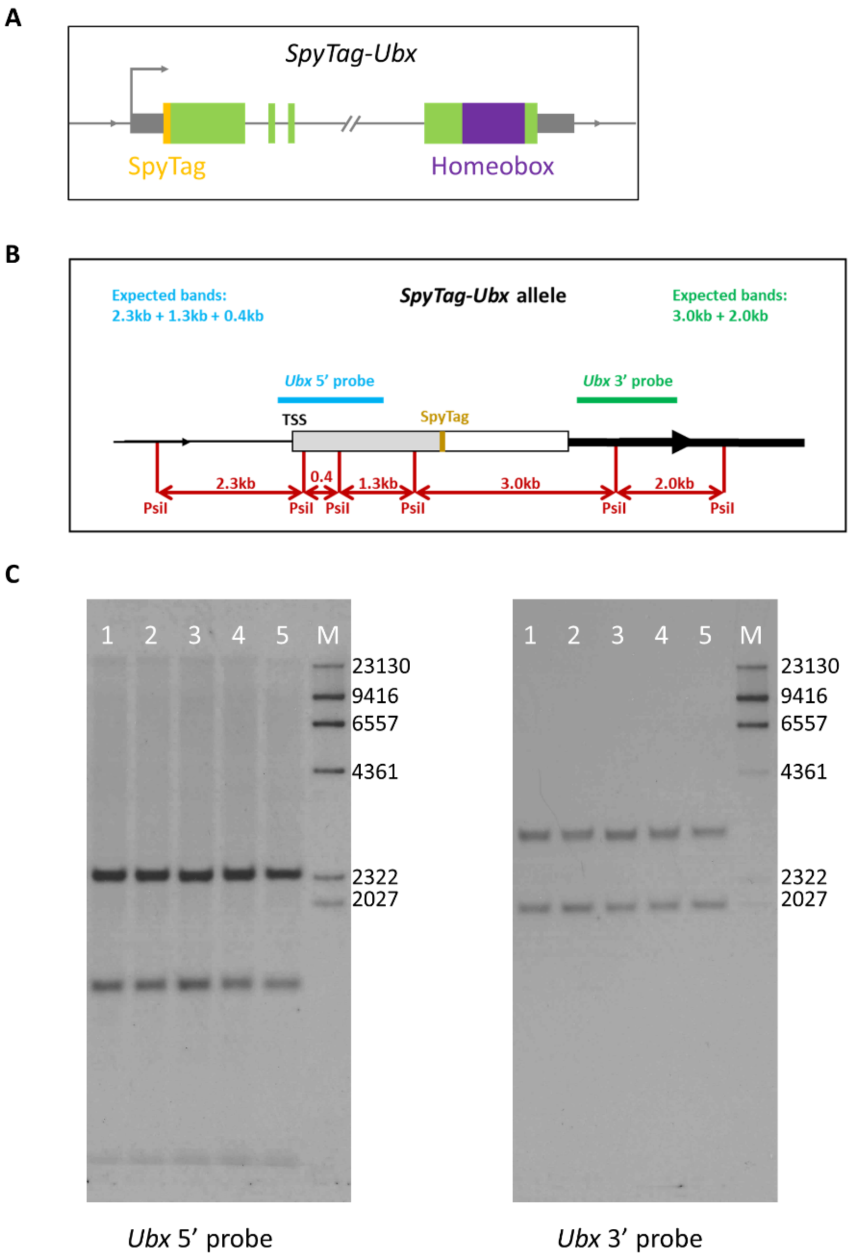


**Fig. S1. Generation and verification of the scarless *SpyTag-Ubx* allele.**

**A.** Schematic of the scarless *SpyTag-Ubx* allele.

**B.** PsiI restriction map of the *SpyTag-Ubx* allele. The sizes of all relevant restriction fragments are shown, and the 5' and 3' Southern blot probe regions are indicated with blue and green bars. The expected Southern blot patterns for each probe are also shown. This schematic is not drawn to scale.

**C.** Southern blot results for five independent *SpyTag-Ubx* alleles. All five alleles gave the expected patterns. Lane M is the marker lane.

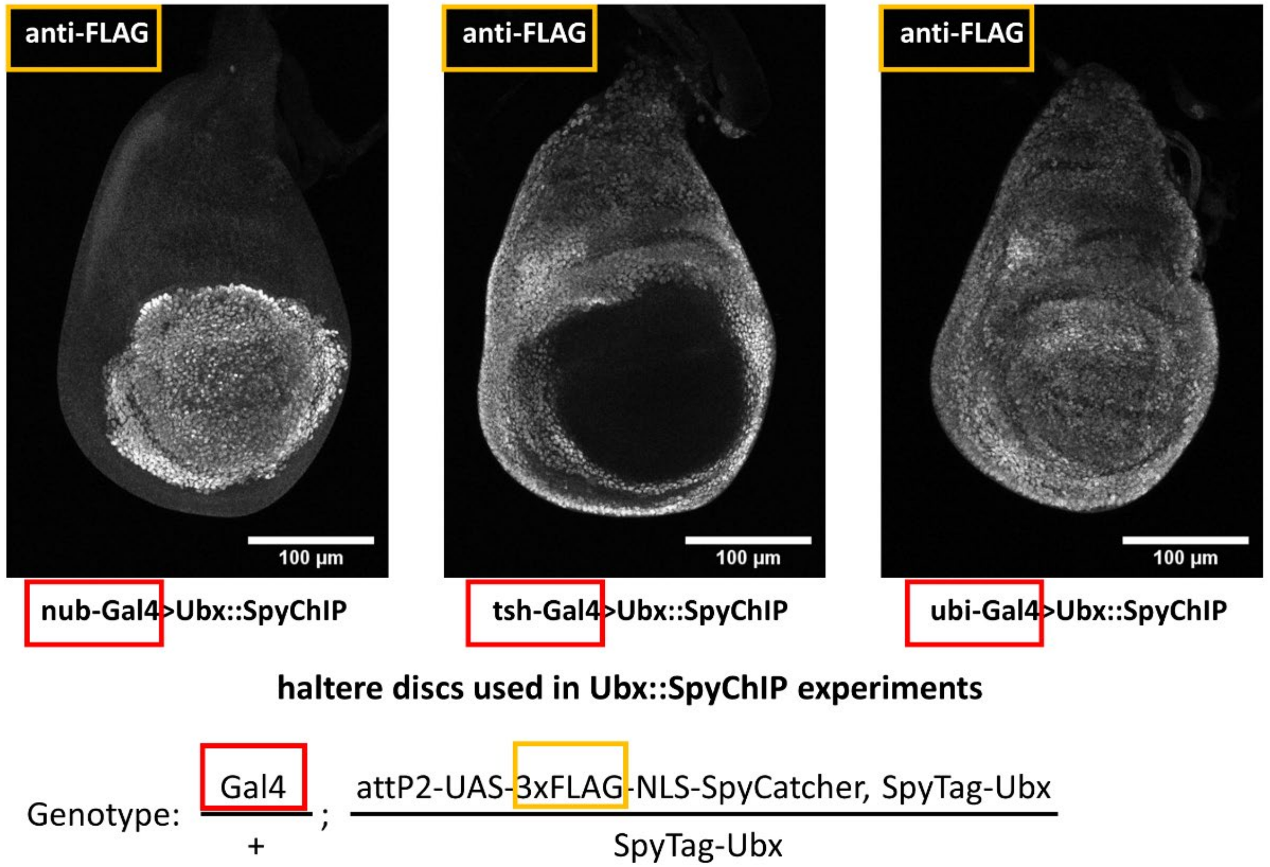


**Fig. S2. Expression of 3xFLAG-SpyCatcher in different domains of *Drosophila* haltere disc.**

Anti-FLAG antibody staining of the haltere discs used in ChIP and western blotting experiments. *tsh-Gal4*, *nub-Gal4* and *ubi-Gal4* all drove *UAS-3xFLAG-SpyCatcher* expression in the expected domains. The detailed genotypes are shown below the images.

**A**

| Ubx ChIP experiment: | Average enrichment (fold) |
|---|---|
| Ubx ChIP with anti-Ubx antibody (Loker *et. al.* 2021) | 2.3 |
| Ubx ChIP with anti-FLAG antibody, using 3xFLAG-Ubx allele | 3.2 |
| ubi-Gal4>Ubx::SpyChIP (anti-FLAG antibody) | 2.1 |
| tsh-Gal4>Ubx::SpyChIP (anti-FLAG antibody) | 2.0 |
| nub-Gal4>Ubx::SpyChIP (anti-FLAG antibody) | 2.1 |

**B**

| Pearson's correlation coefficient | ubi-Gal4>Ubx::SpyChIP (FLAG antibody) | 3xFLAG-Ubx ChIP (FLAG antibody) | anti-Ubx ChIP rep1 (Ubx antibody) |
|---|---|---|---|
| anti-Ubx ChIP rep1 (Ubx antibody) | 0.684 | 0.749 | |
| 3xFLAG-Ubx ChIP (FLAG antibody) | 0.772 | | |
| anti-Ubx ChIP rep2 (Ubx antibody) | | | 0.694 |

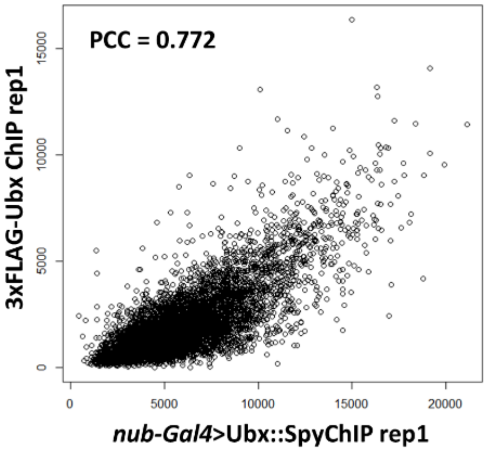


**C**

| Pearson's correlation coefficient | nub-Gal4>Ubx::SpyChIP rep1 (no quenching) | nub-Gal4>Ubx::SpyChIP rep2 (quenching) |
|---|---|---|
| nub-Gal4>Ubx::SpyChIP rep3 (quenching) | 0.891 | 0.893 |
| nub-Gal4>Ubx::SpyChIP rep2 (quenching) | 0.881 | |

**Fig. S3. SpyChIP faithfully captures TF-DNA binding profiles.**

**A.** Average enrichment of sequencing tags in all called peaks relative to a random set of genomic intervals in different Ubx::SpyChIP and whole disc Ubx ChIP experiments.

**B.** Comparisons between *ubi-Gal4*>Ubx::SpyChIP results and two independent whole haltere disc Ubx ChIP results. The Pearson's correlation coefficient values of various pair-wise comparisons are shown. The scatter plot of one comparison is shown on the right. Anti-Ubx ChIP results are from (1), and 3xFLAG-Ubx ChIP was performed in this study with the same anti-FLAG antibody as used in all Ubx::SpyChIP experiments against the 3xFLAG epitope in the scarless *3xFLAG-Ubx* line (2).

**C.** Pair-wise comparisons among different *nub-Gal4*>Ubx::SpyChIP replicates with or without quenching using synthetic SpyTag. The Pearson's correlation coefficient values are shown.

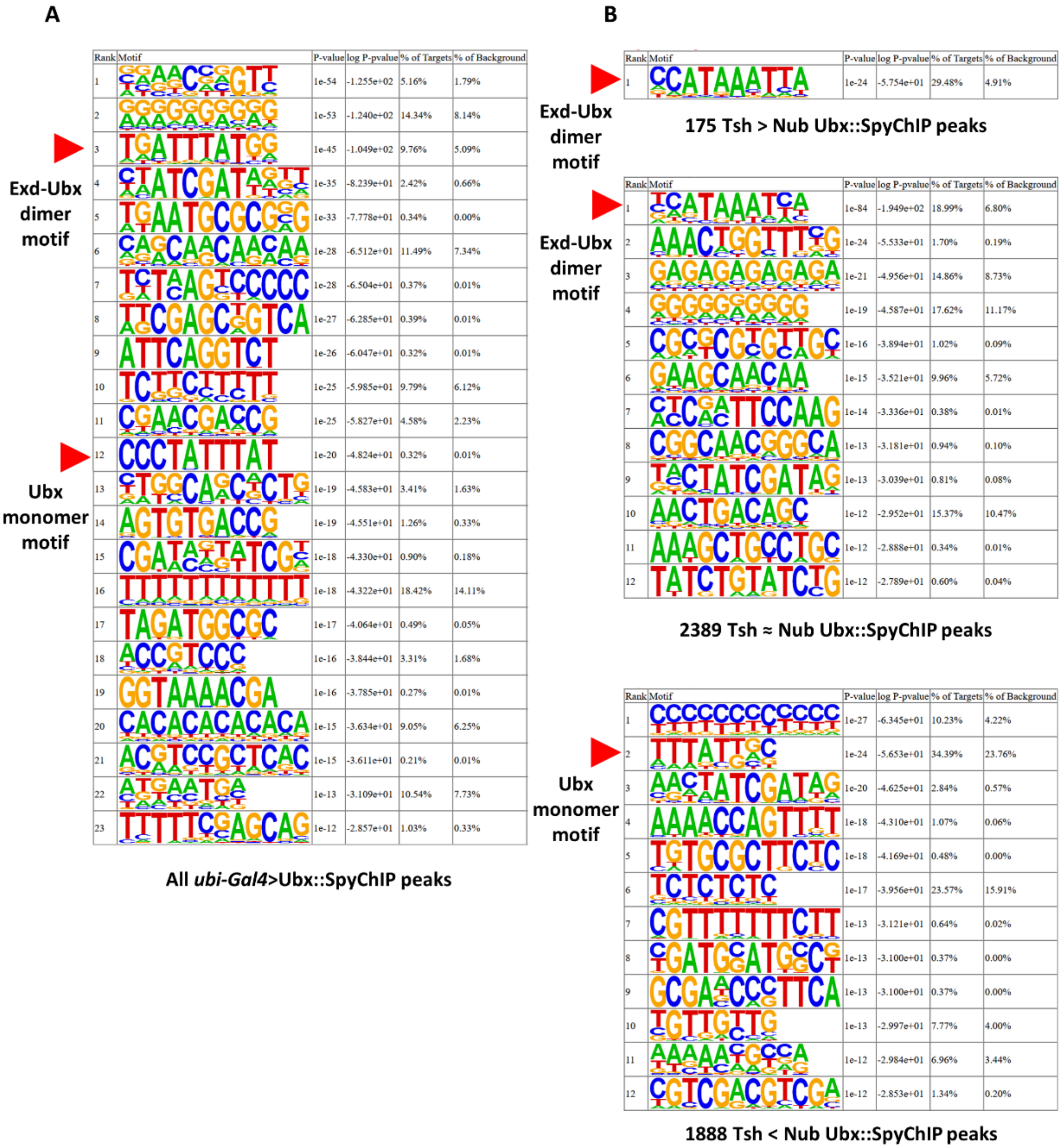


**Fig. S4. All statistically significant motifs enriched in different Ubx::SpyChIP peak sets.**

**A.** Motifs significantly enriched in *ubi-Gal4*>Ubx::SpyChIP peaks. Red arrows point to Ubx monomer and Exd-Ubx dimer motifs. No other Hox motifs are found in these lists.

**B.** Complete lists of significantly enriched motifs discovered through *de novo* motif searches in different classes of Ubx::SpyChIP peaks. Red arrows point to the Hox motifs and are also shown in **Fig. 3B**. No other Hox motifs are found in these lists.

**Extended Methods**

<u>Cloning for generating transgenic flies</u>

The addgene plasmid #26224 was used as the template for the following 2 PCR reactions: the fragment GFPnls was amplified using primers DmGFP-ATG+5' + DmGFP-nls-3', and the fragment GFPnls(ATG-) was amplified using primers DmGFP-ATG-5' + DmGFP-nls-3'. Both fragments were digested with BamHI + EcoRI, and ligated into BamHI + EcoRI digested pBluescript II KS(+), generating constructs PBS-GFPnls and PBS-GFPnls(ATG-) respectively. The oligos V5-STOP-5' and V5-STOP-3' were annealed to form double stranded V5-STOP fragment with EcoRI and HindIII overhangs, which was ligated into EcoRI + HindIII digested PBS-GFPnls and PBS-GFPnls(ATG-), generating PBS-GFPnls-V5 and PBS-GFPnls-V5(ATG-). The ATG-SpyTag fragment was generated by annealing oligos SpyTag-5' and SpyTag-3', and was ligated into NotI + BamHI digested PBS-GFPnls-V5(ATG-). This ligation created the construct PBS-SpyTag-GFPnls-V5.

The oligos 3xFLAG-NLS-1-5' and 3xFLAG-NLS-1-3' were annealed to generate double stranded DNA fragment 3xFLAG-NLS-1, and similarly, oligos 3xFLAG-NLS-2-5' and 3xFLAG-NLS-2-3' were annealed to generate double stranded DNA fragment 3xFLAG-NLS-2. Both fragments were phosphorylated by T4 polynucleotide kinase, and were then ligated into NotI + XbaI digested pBluescript II KS(+), generating the construct PBS-3xFLAG-NLS. The addgene plasmid number 35044 was used as the template to PCR amplify the SpyCatcher fragment using primers SpyCatcher-5' + SpyCatcher-3'. This fragment was digested with XbaI + BamHI, and was ligated into XbaI + BamHI digested pBluescript II KS(+), generating PBS-SpyCatcher construct. The 3xFLAG-NLS fragment was released from PBS-3xFLAG-NLS by digestion with NotI + XbaI, and was ligated into NotI + XbaI digested PBS-SpyCatcher, generating the construct PBS-3xFLAG-NLS-SpyCatcher.

The fragments GFPnls-V5, SpyTag-GFPnls-V5, and 3xFLAG-NLS-SpyCatcher were released from PBS-GFPnls-V5, PBS-SpyTag-GFPnls-V5, and PBS-3xFLAG-NLS-SpyCatcher by digestion with NotI + Acc65I, and subcloned into NotI + Acc65I digested pUASTattB vector, generating pUASTattB-GFPnls-V5, pUASTattB-SpyTag-GFPnls-V5, and pUASTattB-3xFLAG-NLS-SpyCatcher plasmids.

The plasmid PBS-GFPnls (see above) was used as template for the following 2 PCR reactions: the fragment V5-GFPnls was amplified using primers V5-GFP-5' + GFP-3', and the fragment V5-GFPnls-SpyTag was amplified using primers V5-GFP-5' + GFP-SpyTag-3'. Both fragments were digested with KpnI$^{HF}$ + NotI$^{HF}$, and ligated into KpnI$^{HF}$ + NotI$^{HF}$ digested pUASTattB vector, generating pUASTattB-V5-GFPnls and pUASTattB-V5-GFPnls-SpyTag.

All pUASTattB plasmids were integrated into selected attP sites through phiC31 integrase mediated site-specific integration. attP2 was used for all GFP constructs, and the SpyCatcher construct was integrated into both attP2 and attP40.

<u>Details of generating the *SpyTag-Ubx* line</u>

PBS-Ubx-N1, PBS-Ubx-N2 and PBS-Ubx-N3 constructs, as well as the targeting vector and the landing sites, have been described in detail (2). The construct PBS-Ubx-N2 was linearized with DraIII$^{HF}$ and gel purified, which was used as the template for the following 2 PCR reactions: the 0.9kb fragment Ubx-N4-SpyTag was amplified using primers M13 + Ubx-N4-SpyTag-3', and the 1kb SpyTag-Ubx-N5 fragment was amplified with primers SpyTag-Ubx-N5-5' + M13R. These two fragments were both gel purified and used as templated for overlapping PCR using M13 and M13R, which gave the 1.9kb fragment SpyTag-Ubx-N2. This fragment was digested with SacII + KpnI, and then ligated into SacII + KpnI digested pBluescript II KS(+) to generate the construct PBS-SpyTag-Ubx-N2. The Ubx-N1 fragment was released from the construct PBS-Ubx-N1 by digestion with XbaI + EcoRI (+ ScaI$^{HF}$ to facilitate gel cutting), and the SpyTag-Ubx-N2 fragment was released from the construct PBS-SpyTag-Ubx-N2 by EcoRI + HindIII digestion. Both fragments were ligated into XbaI + HindIII digested PBS-Ubx-N3 construct, which generated the construct PBS-SpyTag-Ubx-N. The SpyTag-Ubx-N fragment was released from the PBS-SpyTag-Ubx-N construct by digestion with PacI + AgeI, and was ligated into PacI + AgeI digested pTargeting-RMCE-insulated targeting vector. The Stbl2 cells (Invitrogen 10268019) were used in this step, which generated the final targeting plasmid pTargeting-SpyTag-Ubx-N.

The *SpyTag-Ubx* allele was generated using the same detailed protocol as described in (2). Briefly, the pTargeting-SpyTag-Ubx-N targeting plasmid was injected into the F1 embryos of the cross between vas-int(X) females and males of the *Ubx* landing site line (*Ubx-attP-ubiDsRed-attP*). Successful RMCE transformants were identified by the presence of *w+* and *3xP3-RFP* markers, as well as the concurrent loss of the *ubiDsRed* marker. The I-SceI and I-CreI homing nucleases were then expressed by heat shock, which introduced double stranded DNA breaks that induced homology directed repair of the *Ubx* locus to the scarless *SpyTag-Ubx* allele. The successful repair products were identified by the lost of both *w+* and *3xP3-RFP* markers. All final alleles were fully verified by Southern blotting and sequencing.

Primers used:

| Oligo Name | Oligo Sequence | Note |
|---|---|---|
| DmGFP-ATG+5' | CAGT GGATCC **CAAA** ATGTCCAAAGGTGAAGAACTG | Start codon, **Kozak sequence,** BamHI site |
| DmGFP-ATG-5' | AGTC GGATCC TCCAAAGGTGAAGAACTGTTTAC | BamHI site |
| DmGFP-NLS-3' | CATG GAATTC ACCTCCTGAGCCTCC **TACCTTTCTCTTCTTTTTTGGAGG** ACCTCCACTTCCGCC CTTGTAGAGCTCATCCATGC | EcoRI site, GGSGG linker, **SV40 NLS** |
| V5-Stop-5' | AATTC GGTAAGCCTATCCCTAACCCTCTCCTCGGTCTCGATTCTACG TAATAG A | Stop codons, V5 Tag, HindIII overhang, EcoRI overhang |
| V5-Stop-3' | AGCTT CTATTA CGTAGAATCGAGACCGAGGAGAGGGTTAGGGATAGGCTTACC G | Stop codons, V5 Tag, HindIII overhang, EcoRI overhang |
| SpyTag-5' | GGCCGC **CAAA** ATGGGAGCCCACATCGTGATGGTGGACGCCTACAAGCCGACGA AG GGGGGATCTGGTGGT G | SpyTag, **Kozak sequence,** start codon, GGSGG linker, NotI overhang, BamHI overhang |
| SpyTag-3' | GATCC ACCACCAGATCCCCC CTTCGTCGGCTTGTAGGCGTCCACCATCACGATGTGGGCTCCCAT TTTG GC | SpyTag, **Kozak sequence,** start codon, GGSGG linker, NotI overhang, BamHI overhang |
| 3xFLAG-NLS-1-5' | GGCCGCAAA **ATG**GACTACAAAGACCATGACGGTGATTATAAAGATCATGACAT CGATTAC | NotI overhang, Kozak sequence, **start codon** |

| | | |
|---|---|---|
| 3xFLAG-NLS-1-3' | TTGTAATCGATGTCATGATCTTTATAATCACCGTCATGGTCTTTGT AGTCC**CAT**TTTGC | NotI overhang, Kozak sequence, **start codon,** complementary overhang for 3-frag ligation |
| 3xFLAG-NLS-2-5' | AAGGATGACGATGACAAGCAACATTCTACTCCTCCAAAAAAGAA GAGAAAGGTAGAA T | XbaI overhang, complementary overhang for 3-frag ligation |
| 3xFLAG-NLS-2-3' | CTAGA TTCTACCTTTCTCTTCTTTTTTGGAGGAGTAGAATGTTGCTTGTCA TCGTCATCC | XbaI overhang |
| SpyCatcher-5' | CGTA TCTAGA GGCGCCATGGTTGATACCTT | XbaI site |
| SpyCatcher-3' | CTGA GGATCC TTAAATATGAGCGTCACCTTTAGTTG | BamHI site, stop codon |
| V5-GFP-5' | GACT GCGGCCGCAAA **ATGGGTAAGCCTATCCCTAACCCTCTCCTCGGTCTCGATTCTACG** TCTGCC TCCAAAGGTGAAGAACTGTTTACC | NotI site, Kozak sequence, **V5 tag,** linker |
| GFP-SpyTag-3' | GACT GGTACC TCACTTCGTCGGCTTGTAGGCGTCCACCATCACGATGTGGGCTCC CTTGATATCGAATTCACCTCC | KpnI site, SpyTag + STOP |
| GFP-3' | GACT GGTACC TCA CTTGATATCGAATTCACCTCC | KpnI site, STOP |
| M13 | GTAAAACGACGGCCAGT | -20, 17mer |
| M13R | CAGGAAACAGCTATGAC | -26, 17mer |
| Ubx-N4-SpyTag-3' | CTTCGTCGGCTTGTAGGCGTCCACCATCACGATGTGGGCTCC CATTGCGCTGCTGGCGGTAAGAATC | part of SpyTag-linker, overlapping region, ATG |
| SpyTag-Ubx-N5-5' | GACGCCTACAAGCCGACGAAGTCTGGCGGTGGGGGCTCTGGTG GCGGAGGATCAGGTGGAGGTGGC AACTCGTACTTTGAACAGGCCTC | part of SpyTag-linker, overlapping region |

### References:


1. Loker R, Sanner JE, & Mann RS (2021) Cell-type-specific Hox regulatory strategies orchestrate tissue identity. *Current Biology* 31(19):4246-4255.e4244.
2. Feng S, Lu S, Grueber WB, & Mann RS (2021) Scarless engineering of the Drosophila genome near any site-specific integration site. *Genetics* 217(3).
